# Multiple Sex-Specific Differences in the Regulation of Meiotic Progression in *C. elegans*

**DOI:** 10.1101/2020.03.12.989418

**Authors:** Sara M. Fielder, Rieke Kempfer, William G. Kelly

**Affiliations:** Biology Department, Emory University, Atlanta, Georgia 30322; Program in Genetics and Molecular Biology, Emory University, Atlanta, Georgia 30322; Epigenetic Regulation and Chromatin Architecture Group Berlin Institute for Medical Systems Biology, Max-Delbrück Centre for Molecular Medicine, Hannoversche Str. 28, 10115 Berlin, Germany

**Keywords:** meiosis, checkpoint, sex-specific checkpoint, *C. elegans*

## Abstract

Meiosis is a highly conserved sexual process, yet significant differences exist between males and females in meiotic regulation in many species. Meiotic progression in C. elegans males proceeds more rapidly than female meiosis, suggesting that female meiotic regulation may be more stringent than in males. We have identified multiple differences in the regulation of synapsis, including a difference that suggests the presence of a female-specific meiotic checkpoint that senses the proper initiation of synapsis. This checkpoint is detected by sex differences in the targeting of histone H3 lysine 9 dimethylation (H3K9me2) to unsynapsed chromatin. During oogenic meiosis in hermaphrodites, the failure to initiate synapsis leads to failure to target H3K9me2 enrichment on unsynapsed chromosomes. Loss of the pachytene checkpoint does not reintroduce H3K9me2 enrichment in hermaphrodites, indicating these checkpoints are separable. In contrast, widespread H3K9me2 enrichment occurs as a result of loss of synapsis initiation in both male meiosis and during spermatogenic meiosis in larval XX hermaphrodites. Additionally, male synapsis is insensitive to loss of the dynein motor light chain DLC-1 and to elevated temperatures, whereas female synapsis is prevented by both conditions. We also show that loss of spindle assembly checkpoint proteins, which provide a kinetic barrier to meiotic progression and are required for DLC-1-dependent synapsis phenotypes in hermaphrodites, does not speed up the rate of synapsis in spermatogenic meiosis. These results indicate that meiosis proceeds more rapidly in males because males lack barriers to meiotic progression that are activated by defective synapsis initiation in females.

## INTRODUCTION

Meiotic synapsis of homologous chromosomes is required in both oogenesis and spermatogenesis to prevent sterility and developmental defects caused by aneuploidies in the offspring. *C. elegans* has been developed as an excellent genetic model system for studying meiotic processes, aided by their transparent gonad in which an assembly line of germ cells progressing through sequential stages of meiotic prophase I is readily visible. Germline stem cells, which divide throughout the adult life of each animal, populate a distal mitotic region, and then move proximally and exit mitosis to enter the leptotene and zygotene stages of Meiosis I. In this region, called the “transition zone” (TZ), the chromosomes condense and cluster together at the nuclear periphery, yielding characteristic crescent shaped nuclei (reviewed in (Lui and Colaiacovo 2013)). This organization is mediated by Zinc finger In Meiosis (ZIM) proteins, each of which bind to specific DNA sequences, named “pairing centers”, on one end of one or two chromosomes (described below), and help attach them to each other and to the nuclear envelope (Phillips and Dernburg 2006; Phillips *et al.* 2009). While pairing and homology are being established, lateral elements of the synaptonemal complex (HIM-3, HTP-3, etc) assemble onto chromosomes, and form an axis onto which central element (CE) components (e.g., SYP proteins) assemble (Zetka *et al.* 1999; Couteau and Zetka 2005; Goodyer *et al.* 2008; Kohler *et al.* 2017). The inner nuclear membrane protein ZYG-12 connects chromosomes to the motor complex dynein in the cytoplasm through binding to the outer nuclear membrane protein SUN-1 (Penkner *et al.* 2007; Sato *et al.* 2009). This trans-nuclear membrane connection to cytoplasmic dynein motors is proposed to move chromosomes around the nuclear periphery during their search for their homologs (Penkner *et al.* 2007; Sato *et al.* 2009; Wynne *et al.* 2012; Woglar and Jantsch 2014).

Defects in or depletion of dynein motor components cause a decrease but not complete cessation of movements of chromosome ends around the nuclear periphery (Wynne *et al.* 2012), and pairing takes longer to occur, increasing the TZ length (Sato *et al.* 2009). Less than half of nuclei will have an incorrectly paired chromosome II with a defect in the dynein complex, implying that dynein induced movements aid, but are not completely necessary for correct homolog pairing (Sato *et al.* 2009). As mentioned, in *C. elegans* one end of each chromosome is enriched for genetic elements that comprise pairing centers that aid in homolog identification and pairing (Macqueen *et al.* 2005; Phillips *et al.* 2005; Phillips and Dernburg 2006). ZIM proteins recognize and bind to these elements, and inter-ZIM protein interactions help to match up potential homologous chromosomes (Macqueen *et al.* 2005; Phillips *et al.* 2005; Phillips and Dernburg 2006). Because some ZIM proteins can bind to more than one specific chromosome, the proposed role of the dynein motor is to provide forces to test the strength of interactions within each paired homolog set and pull apart incorrect matches. Once a match has survived this test, SYP proteins (SYP-1, −2, −3, and −4) form a dynamic, physical bridge that “zips” the homologs together starting near ZIM proteins and processing down the length of the condensed chromosomes (Schild-prufert *et al.* 2011; Rog and Dernburg 2015; Rog *et al.* 2017). Depletion of dynein motor components has been reported to cause off-chromatin assembly of SYP proteins into “polycomplexes”, as does deletion of a lateral element such as HTP-3 (Sato *et al.* 2009; Zhang *et al.* 2015; Rog *et al.* 2017). The polycomplexes resulting from deletion of HTP-3 have a structural organization similar to that found in the chromosome-associated synaptonemal complex (Rog *et al.* 2017). Synapsis in hermaphrodites is also inherently temperature sensitive, as exposure to temperatures above 26.5° results in unrecoverable aggregation of SYP proteins into structures that are different than polycomplexes (Bilgir *et al.* 2013; Rog *et al.* 2017). Sequestration of SYP proteins into these polycomplexes and aggregates results in failure of meiosis in these nuclei.

As homologous chromosomes complete synapsis, they disburse within the nucleus and then enter the pachytene stage, where double-stranded breaks initiate and recombination occurs. In *C. elegans*, any chromosomes or chromosome segments that remain unsynapsed, such as the single male X, are targeted for enrichment of histone H3 lysine 9 dimethylation (H3K9me2)(Bean *et al.* 2004). Related processes occur in mice and other organisms, called meiotic silencing of unsynapsed chromosomes (MSUC), with enrichment of repressive epigenetic marks on any chromosomes or segments that fail to completely synapse (Khalil *et al.* 2004; Turner *et al.* 2005; Turner *et al.* 2006)(reviewed in (Turner 2007). Despite the enrichment of the heterochromatin mark, it is unclear whether genes on the enriched chromosomes are actually “silenced”, but the H3K9me2 enrichment has been proposed to help “hide” the unsynapsed X chromosome from the synapsis checkpoint machinery in *C. elegans* males (Checchi and Engebrecht 2011). In this synapsis checkpoint, Pachytene Checkpoint Protein 2 (PCH-2) senses pairing proteins bound to unsynapsed chromatin during pachytene and activate the apoptosis pathway to prevent aneuploid, and thus defective, gametes from developing (Bhalla and Dernburg 2005). Checkpoints in meiosis ensure that important processes have been completed before progressing into subsequent steps, and either prevent progression until that process is completed, or, if they are unable to be completed, sentence cells to apoptosis.

Although the final goal of meiosis –the generation of haploid gametes-is the same in males and females, there are significant differences in basal aspects of meiosis observed between spermatogenesis and oogenesis. An obvious difference is that completion of oogenic meiosis results in a single oocyte, whereas four equal sperm cells are produced from each germ cell in spermatogenesis. Another difference in mammals is that males have more stringent responses to errors in meiosis than females: defects in oocytes that lead to arrest and/or apoptosis in spermatogenesis can by-pass inspection during oogenesis and lead to an increased rate in aneuploidy in mammalian females as compared to mammalian male gametes (Reviewed in (Morelli and Cohen 2005)). However, not all organisms have increased stringency in male meiosis as compared to female meiosis. For example, female *Drosophila* synapse and recombine their chromosomes but neither of these processes occur in males and spermatogenesis proceeds uninterrupted (Morgan 1914). Interestingly, *C. elegans* males complete prophase nearly two and a half to three times faster than females, and it has been speculated that less stringent checkpoints in males may account for this difference (Jaramillo-lambert *et al.* 2007). There is substantial apoptosis during *C. elegans* oogenesis, but apoptosis is inhibited in spermatogenesis, even in cases where meiotic checkpoints are activated (Gumienny *et al.* 1999; Gartner *et al.* 2000). Indeed, male germ cells attempt to repair any damage that would normally set off a checkpoint, yet do not eliminate germ cells that are ultimately unable to repair genomic defects during Prophase I (Jaramillo-lambert *et al.* 2010).

In this study, we identify and examine two additional differences in early meiotic prophase between spermatogenesis and oogenesis. First, we identify an apparent checkpoint, operating only in oogenesis, which senses the initiation of synapsis. This checkpoint is observed when examining the onset of H3K9me2 enrichment on unsynapsed chromatin: if synapsis initiation occurs normally or is only delayed, unsynapsed chromosomes become enriched with this modification. However, if synapsis initiation is not reached in female meiosis, H3K9me2 enrichment is not observed on any chromosomes. The proposed checkpoint appears to engage after pairing is complete but before the onset of synapsis: if synapsis even partially initiates on one or two sets of chromosomes, H3K9me2 enrichment is observed. In contrast, all chromosomes become enriched for H3K9me2 in male mutants lacking any synapsis initiation. Additionally, as previously reported (Sato *et al.* 2009; Zhang *et al.* 2015), when dynein is depleted during leptotene/zygotene in oogenesis, SYP proteins associate into polycomplexes unassociated with chromosomes. However, polycomplexes are not observed in male meiotic cells lacking dynein function. The connection between these processes is not currently understood, but may illustrate a novel and basal sex-specific difference in the stringency of meiotic genome quality control between the two sexes.

## METHODS

### Strains

Strains used in this paper are AV307 (*syp-1*(me17) V/ nT1 (IV;V)), CA258 (*zim-2* (tm574)), N2 Bristol strain, CA1207 (*dhc-1* (ie28[*dhc-1*::degron::GFP]) I), CA1199 (*unc-119*(ed3) III; ieSi38[sun-1p::TIR1::mRuby::sun-1 3’UTR + Cbr-unc-119(+)] IV), wgls227 [syp-2::TY1::EGFP::3xFLAG(92C12) + unc-119(+)]; *syp-2*(ok307) V, *him-5*(e1467), *pch-2*(tm1458) II, *her-1*(e1518), *tra-2*(q276), and *rrf-3(pk1426)* I. The *syp-2:gfp (ck38)V* allele was generated using CRISPR-mediated GFP tagging of the endogenous locus (Fielder and Kelly, manuscript in preparation). All strains were grown at 20° on NGM plates unless noted otherwise noted. Large quantities of synchronized stages were obtained by adding bleach to a final concentration of 10% and incubating for 5-7 minutes to dissolve adults, followed by three washes with M9. Embryos obtained were rocked overnight at room temperature without food to allow hatching and synchronization of L1s. All strains were provided by the CGC, which is funded by NIH Office of Research Infrastructure Programs (P40 OD010440).

### RNA interference

RNAi knockdown of *htp-3* was achieved by feeding animals with HT115 bacterial cells transformed with an *htp-3* RNAi clone (F57C9.5) from Source BioScience (Kamath *et al.* 2003). Knockdown of *dlc-1* was achieved by cloning a 700 base pair long segment from *dlc-1* genomic sequence (see Table 1 for primer pairs) and inserting this segment into the L4440 empty vector multi-cloning site by restriction enzyme cloning, transforming this plasmid into HT115 cells, and feeding the resulting bacteria to animals. Empty vector control RNAi was performed by feeding animals HT115 bacteria transformed with the empty vector RNAi plasmid L4440 (a generous gift from Andrew Fire).

**Table 1.**
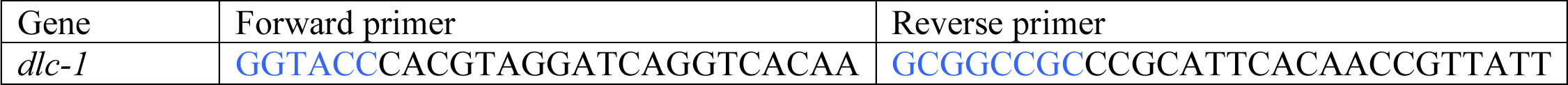
primers used to amplify regions of genes from genomic DNA to insert into L4440 vector for RNAi. Blue letters represent added restriction sequences (such as NotI and KpnI) for ease of addition into the multi-cloning site of the L4440 vector.

For all RNAi knockdown experiments, animals were fed with dsRNA expressing bacteria starting from the L1 larval stage at 20° until 24 hours past the larval L4 stage, and were then shifted to 25° for 24 or 48 hours before examination by immunofluorescence, except where noted differently in Figure 2.

### Immunofluorescence

Immunofluorescence was performed essentially as previously described (Ahn *et al.* 2017), except animals were fixed with 1% final concentration of paraformaldehyde, then slides were frozen with coverslips for less than five minutes on a dry ice block followed by incubation in 100% ethanol at −20° for 2 minutes. Primary antibodies used were goat anti-SYP-1 (1:1500)(Harper *et al.* 2011) guinea pig anti-HTP-3 (1:500)(Macqueen *et al.* 2005), guinea pig anti-ZIM-2 (1:200)(Phillips and Dernburg 2006), rabbit anti-ZIM-1 (1:200)(Phillips and Dernburg 2006), rabbit anti-ZIM-3 (1:200)(Phillips and Dernburg 2006) (all generous gifts from Abby Dernburg), rabbit anti-MDF-1 (1:500), rabbit anti-MDF-2 (1:500)(Essex *et al.* 2009), rabbit anti-BUB-3 (1:500)(Essex *et al.* 2009), (generous gifts from Arshad Desai), and mouse anti-H3K9me2 (1:500) (Abcam ab1220). Secondary antibodies were used at 1:500 dilution: donkey anti-goat (Invitrogen A11055), donkey anti-rabbit (Invitrogen A21207), donkey anti-mouse (Invitrogen A21203), rabbit anti-guinea pig (ThermoFisher PA1-28595).

### Auxin induction

Auxin induced degradation was performed essentially as previously described (Zhang *et al.* 2015). A stock solution of 400mM auxin (Alfa Aesar #A10556) dissolved in ethanol was kept in a light proof tube at 4° for no more than a week. NGM media was prepared and cooled to approximately 50° before adding 1mM final concentration of auxin. Poured plates were kept in tin foil-lined containers to avoid exposing plates to light. OP50 bacteria were seeded onto auxin plates and grown at room temperature in a dark cabinet for 2 days before adding worms that were 24 hours past larval L4 stage for auxin induction. Auxin exposure to worms took place in a dark 20° incubator for 8 hours before analysis by immunofluorescence.

### Hexanediol treatment

Animals treated with RNAi were dissected on poly-L-lysine coated slides in 1x egg buffer + 2.5mg/mL levamisole to immobilize animals. Slides were immediately live imaged for before pictures, and 2x volume of 10% 1,6-hexanediol (Acros Organics/Fisher Scientific 120650010) dissolved in 1x egg buffer for a final concentration of 6.67% was gently pipetted under the coverslip. Images were taken 30 seconds after addition of hexandeiol.

### Data Availability

All non-commercially available reagents are available upon request.

## RESULTS

### H3K9me2 targeting to unsynapsed chromosomes differs between sexes

Dimethylation of histone H3 on lysine 9 (H3K9me2) is normally detectable at low levels by immunofluorescence in *C. elegans* pachytene stage germ cells, and is largely confined to the ends of the chromosomes (Kelly *et al.* 2002; Reuben and Lin 2002) (Fig. 1). However, this modification becomes enriched when one or more sets of homologs fail to synapse, and is also enriched at unsynapsed chromosomal segments (Kelly *et al.* 2002; Bean *et al.* 2004). Examples include the asynapsed lone X chromosome in males (Bean *et al.* 2004) (Fig. 1), or the two unsynapsed chromosome V’s in *zim-*2 (Fig. 1). Additionally, *zim-1* mutant hermaphrodites that cannot pair or synapse multiple chromosome pairs (LG II and III) display as many as four chromosomes with high enrichment for H3K9me2 (data not shown). Thus when multiple chromosomes remain unsynapsed, they are targeted for H3K9me2 enrichment in hermaphrodite germ cells. Importantly, in these cases the remaining chromosomes still engage in normal synapsis and are not enriched in H3K9me2. We also investigated mutants that have a complete synapsis defect in all chromosomes, such as occurs in *syp-1* mutants. Surprisingly, in the complete absence of synapsis no enrichment of H3K9me2 was observed on any chromosomes in the pachytene region of *syp-1* female germ cells (adult hermaphrodite germ cells will be referred to as female germ cells for clarity; Fig. 1). The absence of H3K9me2 enrichment was also observed in other animals with complete synapsis defects, such as *syp-2* and *him-3* mutants; furthermore, the lack of H3K9me2 enrichment in chromatin was not due to dispersal of a limited signal, as no increase in H3K9me2 above background was observed (data not shown). As previously shown, mutants with complete asynapsis also display an extended region of condensed, crescent-shaped chromatin that is normally restricted to the transition zone, indicating that meiotic progression has stalled in these nuclei (MacQueen *et al.* 2002).

**Figure 1.**
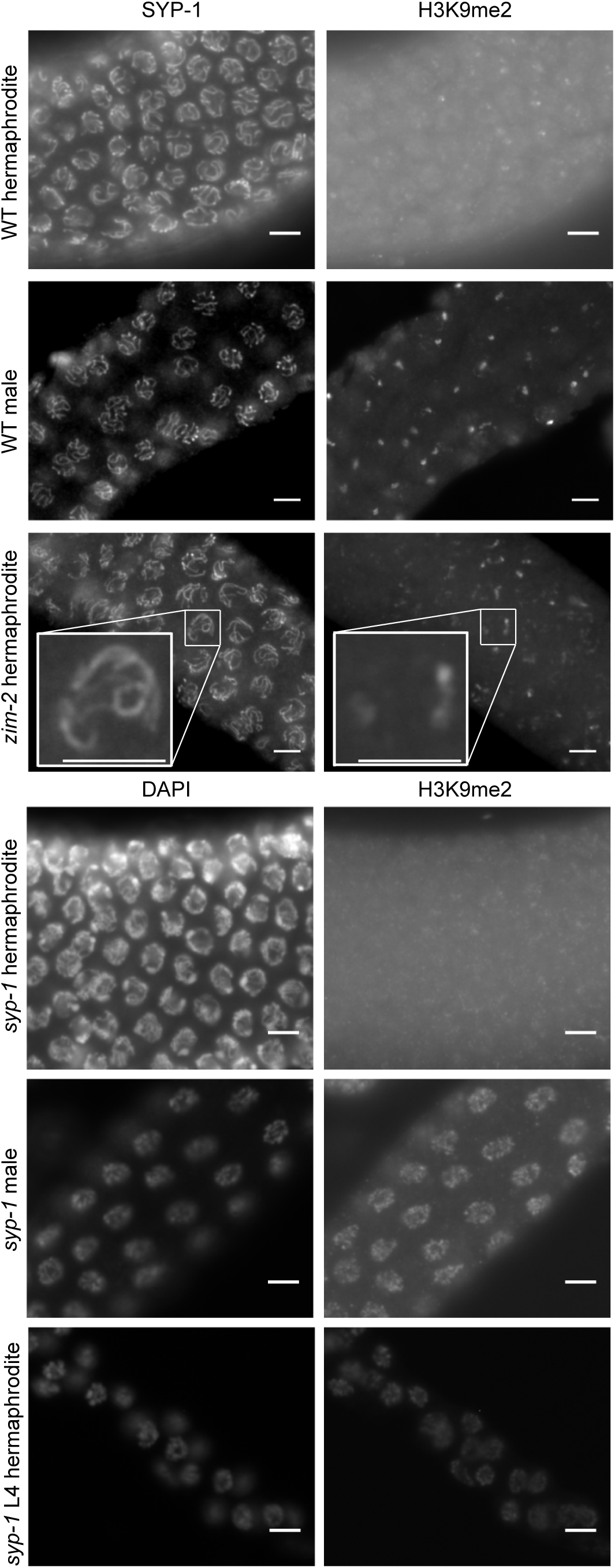
Germline sex-specific H3K9me2 enrichment in response to meiotic synapsis defects. Whole mount fixed gonads were probed with anti-SYP-1 and anti-H3K9me2 (top 3 rows) and/or counterstained with DAPI (bottom 3 rows, with meiosis progressing from left to right in all images). Chromatin in WT female synapsis have low levels of H3K9me2 (top row), while male germ cells have the unsynapsed, single X chromosome that is highly enriched for H3K9me2 (second row). *zim-2* hermaphrodites have defective pairing and synapsis of LG Vs, and display two chromosomes that accumulate high levels of H3K9me2 (third row, insets). *syp-1* female germ cells cannot initiate synapsis and have low levels of H3K9me2 (fourth row), whereas *syp-1* male germ cells exhibit widespread H3K9me2 (fifth row). *syp-1* L4 hermaphrodites undergoing spermatogenesis also display widespread H3K9me2 (bottom row). Scale bars 5 μm.

Unexpectedly, in contrast to what we observed in female germ cells, male *syp-1* mutants displayed widespread enrichment of H3K9me2 in germ cell chromatin on all chromosomes in pachytene nuclei (Fig. 1). The enrichment of H3K9me2 on all chromosomes in male germ cells further indicates that the absence of enrichment in female germ cells is not due to a more diffuse or diluted enrichment due to an increase in unsynapsed chromatin substrate. The difference in responses to the loss of synaptonemal components in males versus hermaphrodites is due to germline sex. Furthermore, *syp-1* larval L4 hermaphrodites, which undergo spermatogenesis before switching to oogenesis at the L4/adult molt, display widespread enrichment of H3K9me2 (Fig. 1). These results indicate that there is a differential regulation of H3K9me2 targeting to unsynapsed chromosomes during meiosis in male versus female germlines.

In WT hermaphrodites, a widespread enrichment of H3K9me2 on chromatin normally begins in diplotene (Kelly *et al.* 2002; Bessler *et al.* 2010) (Fig. S1). This enrichment coincides temporally with the beginning of removal of SYP proteins in preparation for chromosome separation. This is also observed in *syp-1* hermaphrodites: at the bend of the gonad arm, when diplotene normally occurs, H3K9me2 enrichment is observed (Fig. S1). The H3K9me2 enrichment at the end of pachytene further implies that the absence of this mark earlier in meiotic progression is not due to a dispersal of the signal, nor to a defect in methyltransferase activity. These results imply that female germ cells have separable modes of H3K9me2 regulation: one that requires the initiation of synapsis and is absent in SC mutants, and one that occurs in diplotene that is unaffected by defective synapsis initiation.

### PCH-2 is not required for adult hermaphrodite-specific prevention of H3K9me2 enrichment with disrupted synapsis

PCH-2, a homolog of yeast PCH2, is required for the selective apoptosis of cells with defective synapsis during hermaphrodite meiosis in *C. elegans* and is proposed to regulate the “pachytene checkpoint” in *C. elegans* (Bhalla and Dernburg 2005). PCH-2 is expressed before the onset of meiosis, and is required to restrain meiotic progression kinetics, with meiosis proceeding more rapidly in *pch-*2 mutants. PCH-2 is thus thought to provide a “kinetic barrier” to ensure meiotic quality control (Deshong *et al.* 2014). In contrast to hermaphrodites, defects in *C. elegans* male synapsis do not induce apoptosis, and meiosis proceeds with faster kinetics in males versus hermaphrodites, suggesting that PCH-2’s activity may not be required in males, and may be involved in the hermaphrodite specific regulation of H3K9 methylation (Jaramillo-Lambert *et al.* 2007; Deshong *et al.* 2014). Disabling a PCH-2 dependent checkpoint in hermaphrodites could thus allow the accumulation of H3K9me2 to occur in the complete absence of synapsis. We therefore knocked down *syp-*2 by *GFP(RNAi)* in a line expressing a CRISPR-generated *syp-2:gfp* in the presence or absence of the *pch-2 (tm1458)* mutation. No increased H3K9me2 enrichment was observed in the *pch-2(tm1458); syp-2:gfp(ck38)* female germ cells relative to those in *syp-2:gfp(ck38)*, despite a clear disruption of synapsis by GFP RNAi (Fig. 2). With knockdown of HTP-3 in *pch-2*, there is occasional widespread enrichment of H3K9me2 in female germ cells, but this is mirrored by occasional widespread enrichment of H3K9me2 in female germ cells with knockdown of HTP-3 alone (Fig. S2). Therefore, the pachytene checkpoint that is regulated by PCH-2 does not appear to be involved in the prevention of H3K9me2 accumulation in hermaphrodite mutants unable to initiate synapsis. PCH-2 loading onto the SC is only observed after synapsis has begun, thus its activity may be required after the initiation of synapsis and thus later than the sex-specific process we observe (Deshong *et al.* 2014).

**Figure 2.**
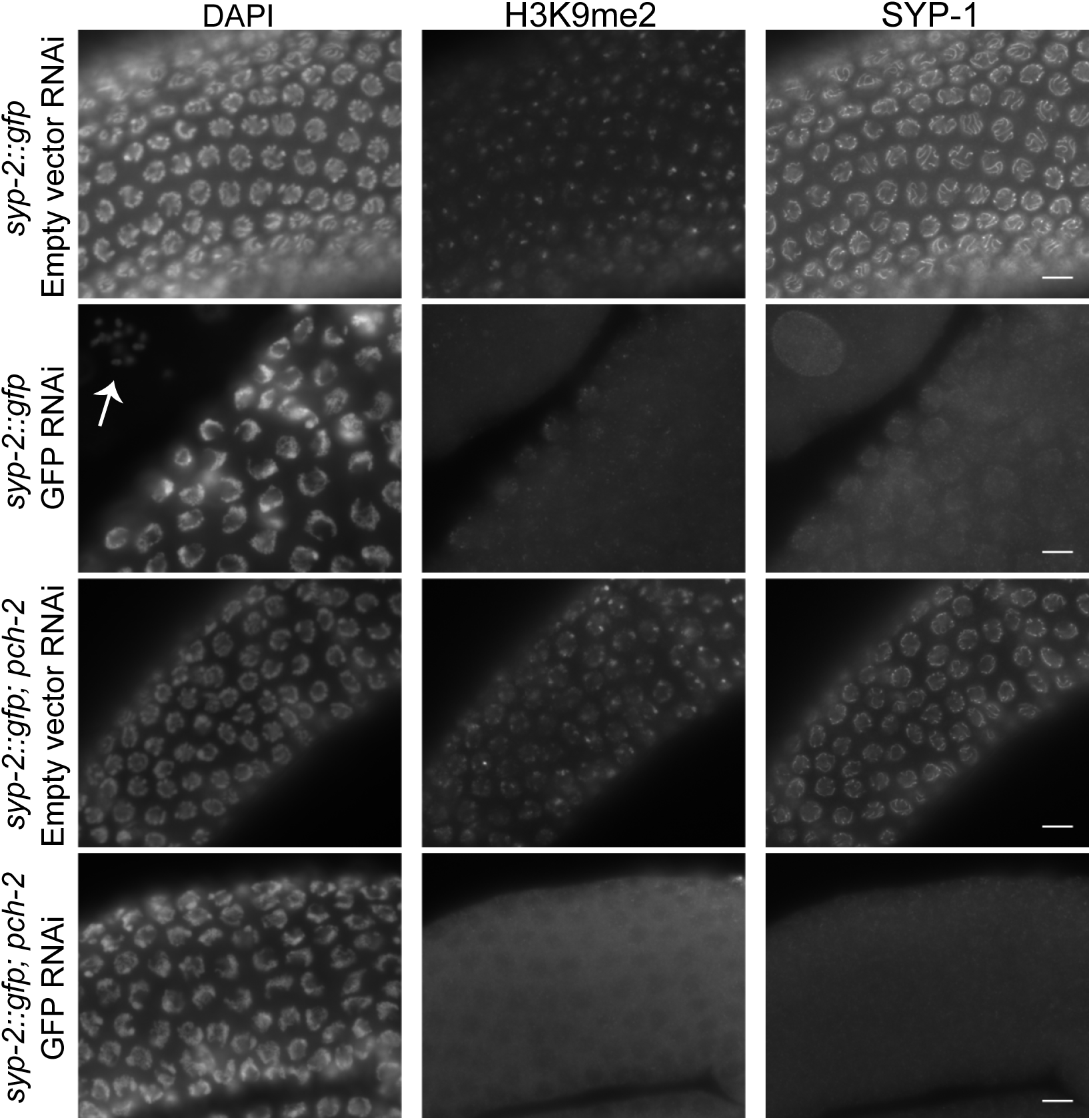
*pch-2* hermaphrodites do not have widespread enrichment of H3K9me2 with absence of synapsis. Whole mount dissected fixed *syp-2::gfp(ck38)* or *syp-2::gfp(ck38); pch-2* gonads probed with anti-H3K9me2, anti-SYP-1, and DAPI. The tagged SYP-2::GFP line gives H3K9me2 enrichment around the telomeres (top row), while knockdown of SYP-2::GFP eliminates much of H3K9me2 (second row). Knockdown of SYP-2::GFP results in increase of DAPI bodies in oocytes (arrow). Deletion of *pch-2* does not result in a change in H3K9me2 patterning in empty vector control RNAi (third row). Deletion of *pch-2* also has low H3K9me2 when SYP-2::GFP is knocked down (bottom row).

### Initiation of synapsis is required for H3K9me2 enrichment of unsynapsed chromosomes in adult hermaphrodite meiosis

In some *htp-3(RNAi)* hermaphrodites we noticed that a few nuclei had small, faint SYP-1 tracks between one or two chromosomes, with the rest of SYP-1 in foci away from chromatin (Fig. 3, arrows). These nuclei had at least partially initiated synapsis, and also displayed enrichment of H3K9me2 on other chromosomes (Fig. 3, arrows). This contrasted with other nuclei in the same gonad that had no or extremely little synapsis and little H3K9me2 enrichment (Fig. 3, arrowheads), We further investigated the temporal dynamics of H3K9me2 targeting to unsynapsed chromatin by utilizing RNAi knockdown of dynein motor light chain 1 (DLC-1) in a temperature shift experiment. *dlc-1(RNAi*) animals grown at 25° form foci of synapsis proteins instead of normal SC tracks between chromosomes (Sato *et al.* 2009). Nuclei that entered meiosis during the period of temperature shift showed SYP-1 foci and low H3K9me2 enrichment (Fig. 3, arrowheads), as was observed in *syp-1* and other mutants with complete synapsis defects. In contrast, more proximal nuclei showed some SYP-1 loading onto chromatin, indicating these nuclei had partially entered meiosis at the onset of the temperature shift (Fig. 3, arrows). These proximal nuclei had very short SYP-1 tracks localized to one or two chromosomes, yet the rest of the chromosomes that lacked SYP-1 were highly enriched for H3K9me2, (Fig. 3, arrows). These results are consistent with a requirement for some level of synapsis initiation to occur before H3K9me2 enrichment on unsynapsed chromosomes can be triggered.

**Figure 3.**
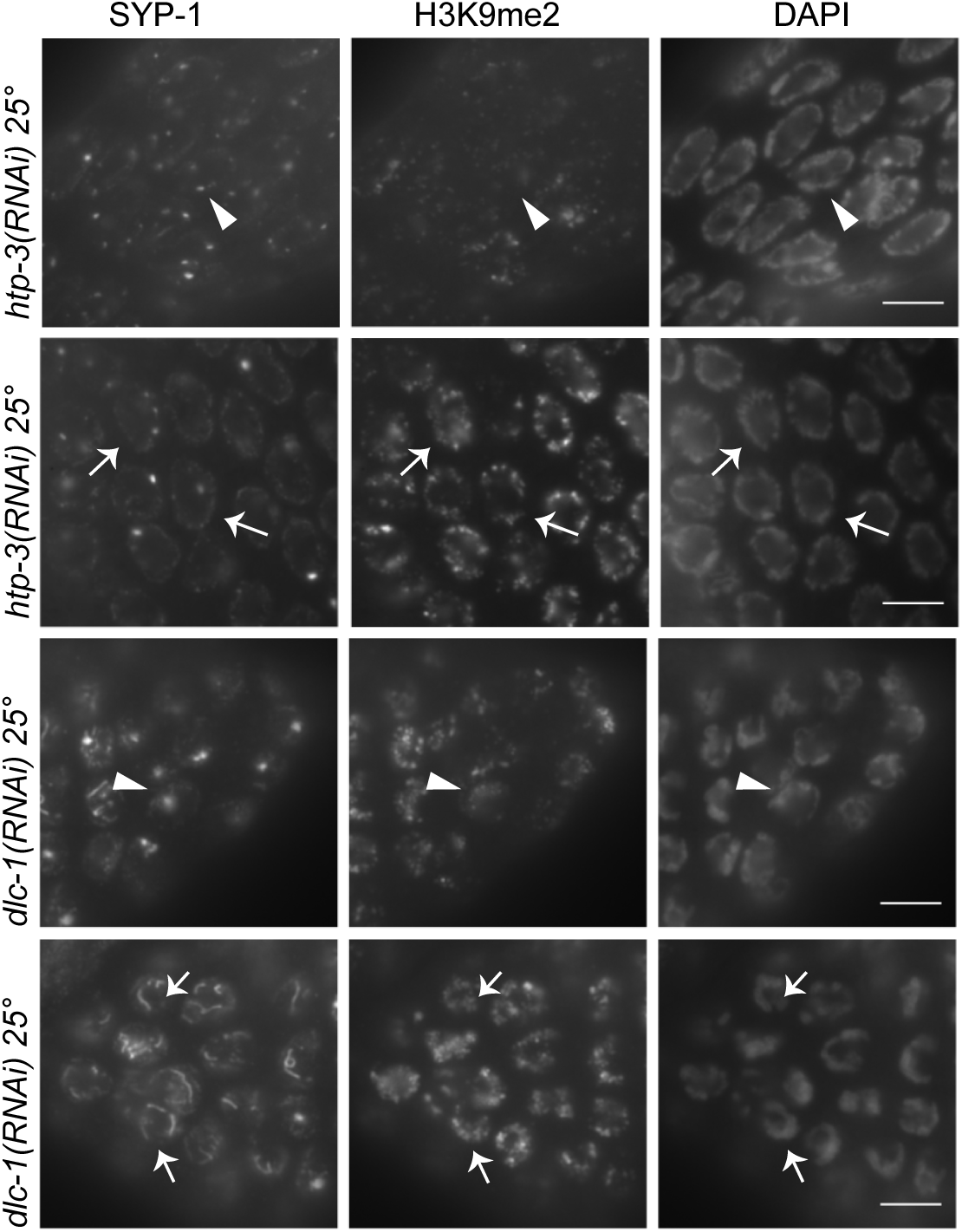
Partial initiation of synapsis is sufficient to trigger widespread H3K9me2 enrichment in female germ cells. Whole mount dissected fixed gonads were probed with anti-SYP-1, anti-H3K9me2, and DAPI. In *htp-3(RNAi)* animals, if SYP-1 proteins are mostly confined to foci (top row, arrowheads), there is little H3K9me2 present. However, in some cases where a small amount of SYP-1 protein has localized to chromatin in *htp-3(RNAi)* animals, there is widespread H3K9me2 enrichment (second row, arrows). Similarly, in *dlc-1(RNAi)* animals, if a nucleus has SYP-1 present only in foci away from chromatin, there is low H3K9me2 (third row, arrowheads), while nuclei that have some, but incomplete synapsis have enrichment of H3K9me2 (bottom row, arrows). Scale bar 5 μm.

### Dynein is not required for males to correctly pair and synapse chromosomes

Dynein motors have been proposed to play an essential role in testing for proper pairing and the subsequent initiation of synapsis (Sato *et al.* 2009). Interestingly, as opposed to hermaphrodites exposed to RNAi in parallel, *dlc-1(RNAi)* males did not exhibit SYP-1 foci, and instead displayed grossly normal synapsis (Fig. 4). In contrast, *htp-3(RNAi)* animals form SYP-1 foci in the germlines of both sexes (Fig. 4), indicating this result is not a difference in RNAi efficiency in the two sexes. To address if the *dlc-1(RNAi)* phenotype is a germline sex or chromosome count specific phenotype, we also tested sex determination mutants: *her-1(e1518)* XO females, and *tra-2(q276)* XX males. SYP-1 foci were observed in *her-1 dlc-1(RNAi)* XO females (Fig. 4), while *tra-2 dlc-1(RNAi)* XX males displayed grossly normal synapsis (Fig. 4). These results suggest that male germ cells, whether of karyotype XO or XX, can initiate synapsis despite a loss of DLC-1.

**Figure 4.**
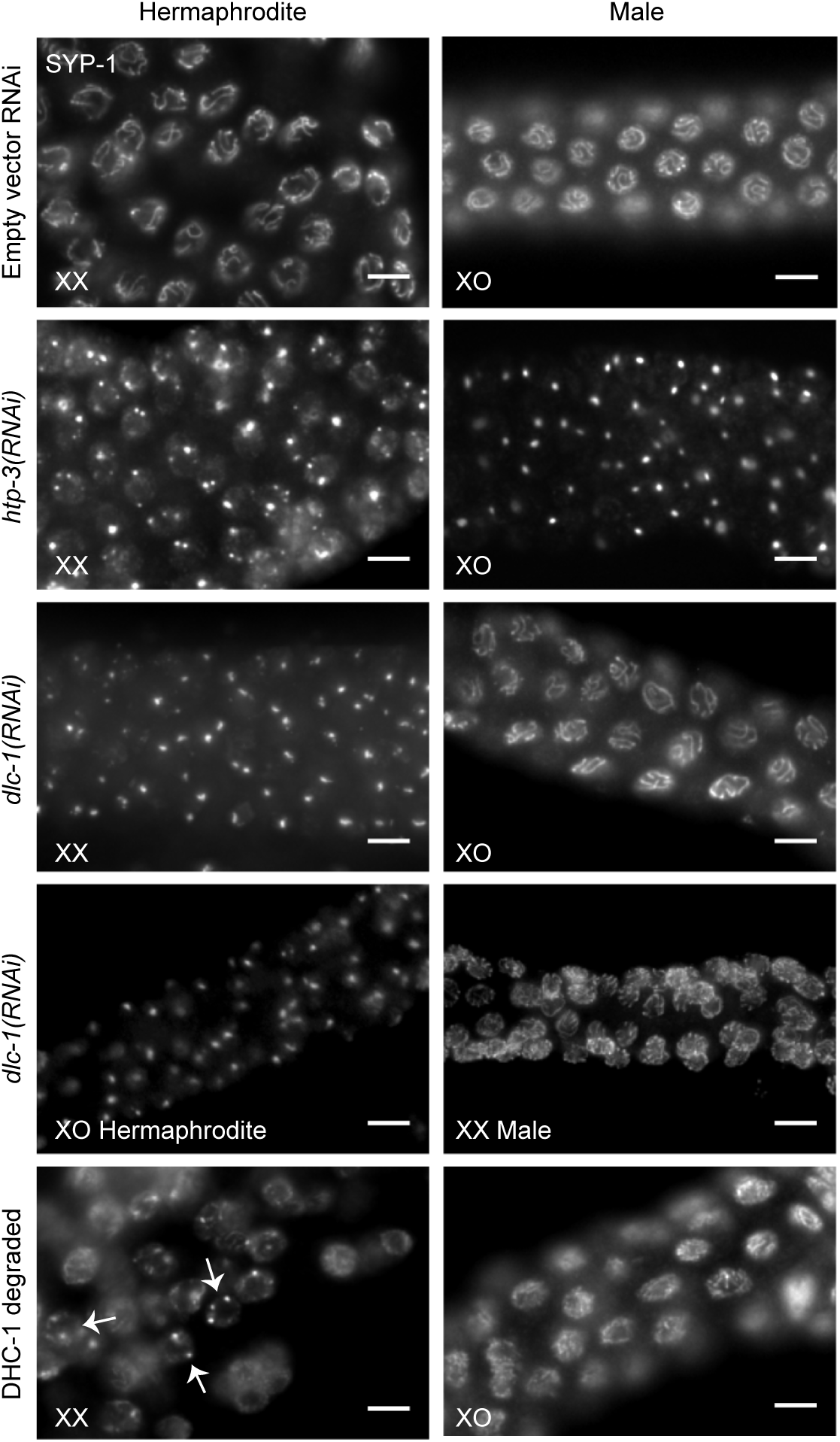
Dynein depletion dramatically affects adult hermaphrodites, but not male synapsis. Whole mount dissected fixed gonads probed with anti SYP-1 are show, with meiotic progression from left to right. WT empty vector RNAi XX hermaphrodites and XO males normally show SYP-1 tracks in between chromosomes (top row). Both *htp-3(RNAi)* XX hermaphrodites and XO males have SYP-1 foci formed away from chromatin (second row). *dlc-1(RNAi)* XX hermaphrodites also form SYP-1 foci away from chromatin, however, *dlc-1(RNAi)* XO males have grossly normal SYP-1 tracks (third row). Depleting DLC-1 in sex determination mutant *her-1(e1518)* XO females display SYP-1 foci, while *tra-2(q278)* XX males have grossly normal synapsis like their male counterparts (fourth row). *dhc-1(ie28)* I; ieSi38 IV animals exposed to 1mM auxin for eight hours at 20° to degrade DHC-1 resulted in XX hermaphrodites that had incidences of SYP-1 foci (arrows), while all XO males analyzed had grossly normal SYP-1 tracks (bottom row). Scale bar 5 μm.

To further investigate the apparent lack of a requirement for dynein in male meiosis, we next investigated whether males require dynein heavy chain (DHC-1), another dynein complex protein, for synapsis. We utilized the auxin inducible degradation (AID) system targeting dynein heavy chain, DHC-1, adapted for use in *C. elegans* by the Dernburg lab (Zhang *et al.* 2015). SYP-1 foci were observed in female germ cells after 8 hours of auxin-induced degradation (Fig. 4). In contrast, males grown on the same plates displayed grossly normal tracks of SYP-1 proteins (Fig. 4) instead of the foci observed in their hermaphrodite counterparts, further indicating that males do not need dynein in order to synapse their chromosomes.

As mentioned, dynein has been proposed to play a role in ensuring the correct pairing of homologs prior to synapsis onset. As dynein is not required for male synapsis, we next investigated whether these males were still able to correctly pair their chromosomes. It has been previously shown that hermaphrodites with depleted dynein have a pairing defect: by the end of meiosis, only 60% of hermaphrodites with depleted dynein correctly pair and synapse chromosome V (Sato *et al.* 2009). We performed immunofluorescence probing ZIM-2 on *dlc-1(RNAi)* males that had been grown at 25° for 24 hours. The region showing ZIM-2 foci was distributed into four zones (Fig. 5A) and pairing status of chromosome V was measured in each nucleus (Fig. 5B). These males have statistically the same rates of pairing as their empty vector RNAi counterparts at all four zones analyzed (Fig. 5C), indicating that males do not need *dlc-1* in order to correctly pair their chromosomes. Further, as pairing can still occur in both *dlc-1(RNAi)* female and male germ cells, but H3K9me2 is not enriched in *dlc-1(RNAi)* hermaphrodites, this indicates that H3K9me2 enrichment of unsynapsed chromosomes normally is triggered after pairing has completed (Sato *et al.* 2009).

**Figure 5.**
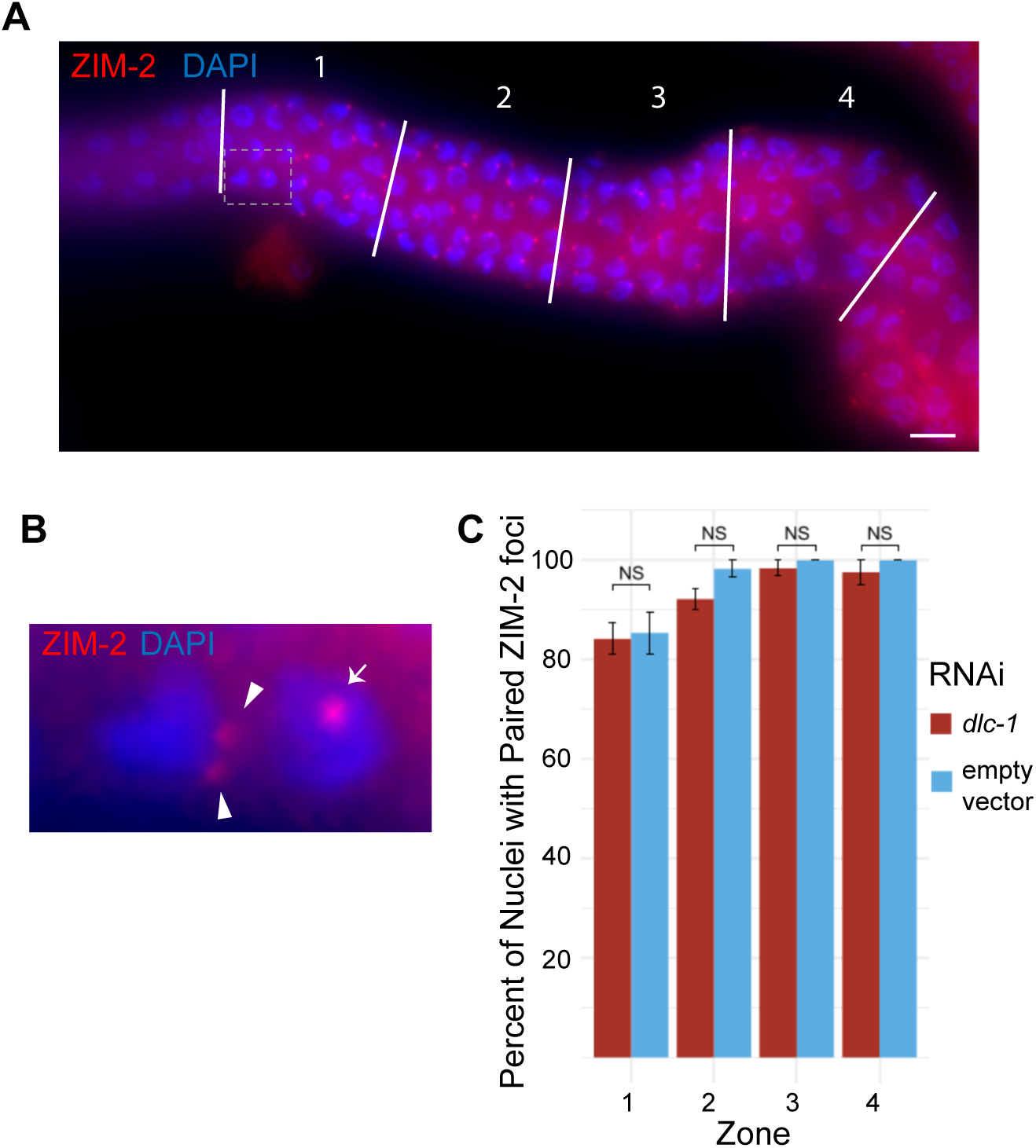
Pairing rates remain unchanged in *dlc-1(RNAi)* males. (A) Whole mount fixed male gonad treated with empty vector RNAi and probed with anti ZIM-2 (red) and DAPI (blue), shown as an example of how the region containing ZIM-2 foci of each gonad was measured, then divided into four equal length zones and each nucleus scored for pairing status of ZIM-2. Dotted box represents portion blown up in panel B. (B) Zoom in of a two nuclei from zone 1, with a nucleus with unpaired, two ZIM-2 foci (arrowheads), and a nucleus with paired, one ZIM-2 focus (arrow) chromosome V. Meiotic progression is depicted from left to right. Scale bar 5 μm. (C) Knockdown of *dlc-1* in N2 males does not change the pairing rates as compared to empty vector RNAi treated N2 control in males. No significance based on in any of the four zones was found using student’s T test. Error bars represent SEM.

### Male synapsis is not as sensitive to heat as hermaphrodite synapsis

As we identified several differences between males and hermaphrodites related to initiation of synapsis, we decided to investigate if the synaptonemal complex formed in males has similar characteristics as that formed in hermaphrodites. Previous electron-microscopic analyses revealed that the foci formed in *htp-*3 mutants have a similar structure, albeit unassociated with chromosomes, to synaptonemal complexes on synapsed chromosomes (Rog *et al.* 2017). They additionally showed that exposure to amphiphilic alcohols, such as 1,6-hexanediol, readily disperses both *htp-3* mutant foci as well as the correctly assembled synaptonemal complex (Rog *et al.* 2017). However, foci formed at higher temperatures (above 26.5°) do not dissolve when exposed to 1,6-hexanediol, implying a change in protein structure (Bilgir *et al.* 2013; Rog *et al.* 2017). Unexpectedly, male synaptonemal complexes are resistant to elevated temperatures, even to 27.5°, and these SYP tracks observed at elevated temperatures still dissolved with exposure to 1,6-hexandiol, similar to hermaphrodite synaptonemal complexes formed under standard growing temperatures (Fig. 6) (Rog *et al.* 2017). Additionally, larval stage L4 hermaphrodite germ cells, which produce sperm, also do not display SYP foci when raised at 27.5° and the SC formed was also readily dispersed by 1,6-hexandiol (Fig. 6). These results suggest that although there is not an obvious structural difference in the SC in male and female germ cells once formed, the formation of the SC appears to have an inherent temperature sensitivity in female meiosis that is not observed in male germlines.

**Figure 6.**
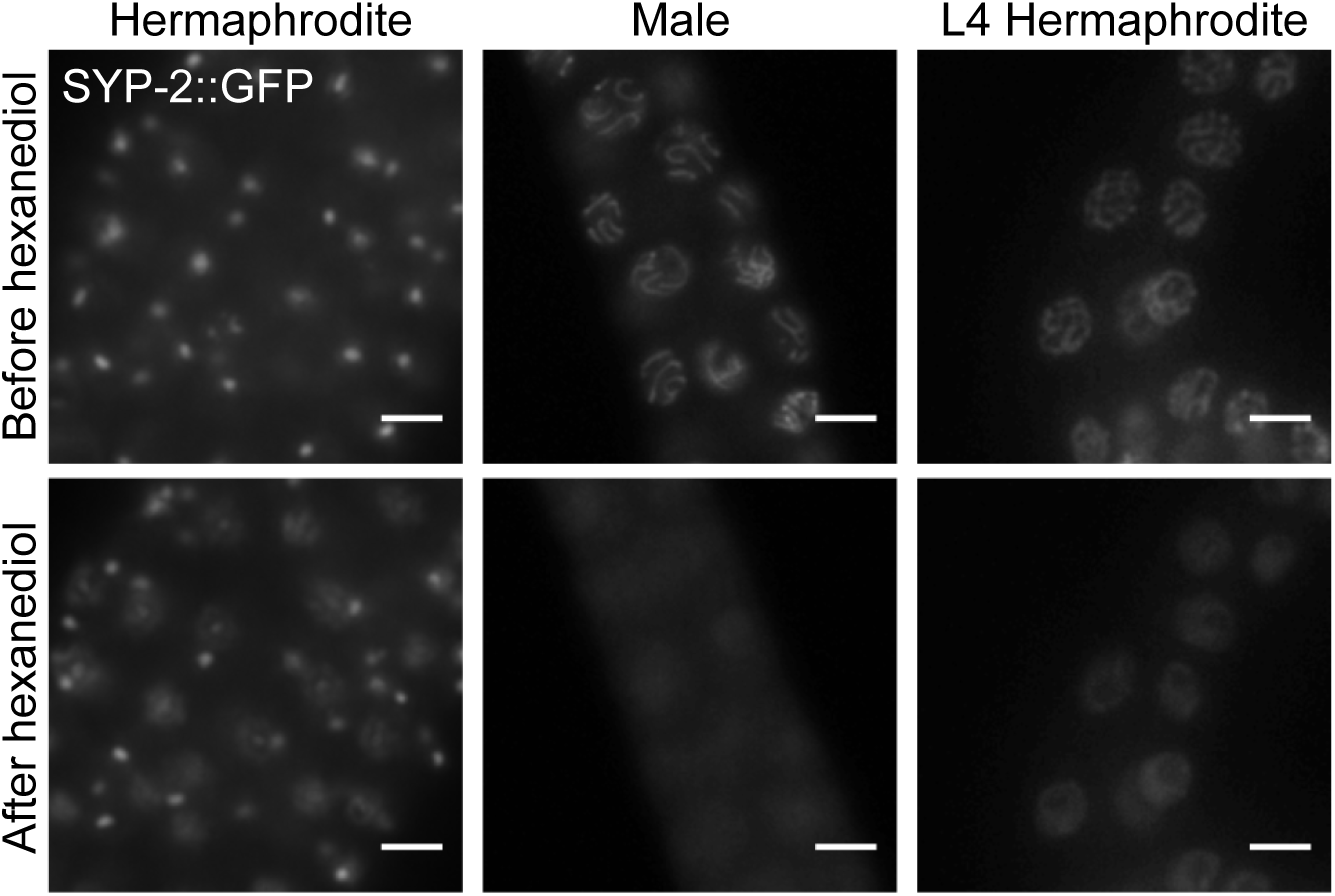
Synaptonemal complex stability is unaffected by increased temperature in male germ cells. Live imaging of extruded gonads of wgls227 (*syp-2::gfp); syp-2(ok307)* V animals. Animals were raised at 27.5° for 24 hours before imaging, starting incubation at larval stage L4 for hermaphrodite and males. Growth at high temperature results in SYP-2 foci in adult hermaphrodites, but males and L4 hermaphrodites, which undergo spermatogenesis, display grossly normal SYP-2 assembly between chromosomes (top row). 30 seconds after addition of 1,6-hexanediol, hermaphrodite SYP-2 foci do not dissolve, but both male and L4 hermaphrodite SC dissociates off of chromosomes to produce a haze around the nucleus of SYP-2. Scale bar 5 μm.

### Spindle assembly checkpoint proteins in males

Male meiosis in *C. elegans* has been shown to progress more rapidly than hermaphrodite meiosis (Jaramillo-lambert *et al.* 2007). In support of this, we noticed that males also appear to complete pairing faster than in hermaphrodites. Empty vector control RNAi males show 98% pairing in the second zone in which ZIM-2 appears (Fig 5C). By comparison, WT adult hermaphrodites do not reach 90% pairing until zone three (Sato *et al.* 2009). Interestingly, it has been shown that meiotic progression speeds up, and the SYP foci phenotype in *dlc-1(RNAi)* hermaphrodites is suppressed in spindle assembly checkpoint (SAC) protein mutants. SYP-1 proteins do not form foci and synapsis is more rapid but appears to proceed normally in these mutants (*mdf-1*, *mdf-2*, and *bub-3*) when combined with *dlc-1(RNAi)* (Bohr *et al.* 2015). Expression data from public datasets also suggested that these proteins are expressed at lower levels in males than in adult hermaphrodites (Gerstein *et al.* 2010). To investigate this, we probed male and adult hermaphrodite germ cells by immunofluorescence using an anti-MDF-2 antibody (Fig. 7)(Essex *et al.* 2009). All three proteins were observed in both male and female germlines (Fig. 7 and data not shown), however, we observed a noticeable difference in MDF-2 localization in male versus female germ cells. Whereas MDF-2 is enriched at the nuclear periphery in and before the transition zone in hermaphrodites (Fig. 7), its distribution is strikingly more diffuse in male transition zone nuclei (Fig. 7). This could imply that MDF-2’s function in males differs from its function in female germ cells and may not be required for normal meiotic progression in males.

**Figure 7.**
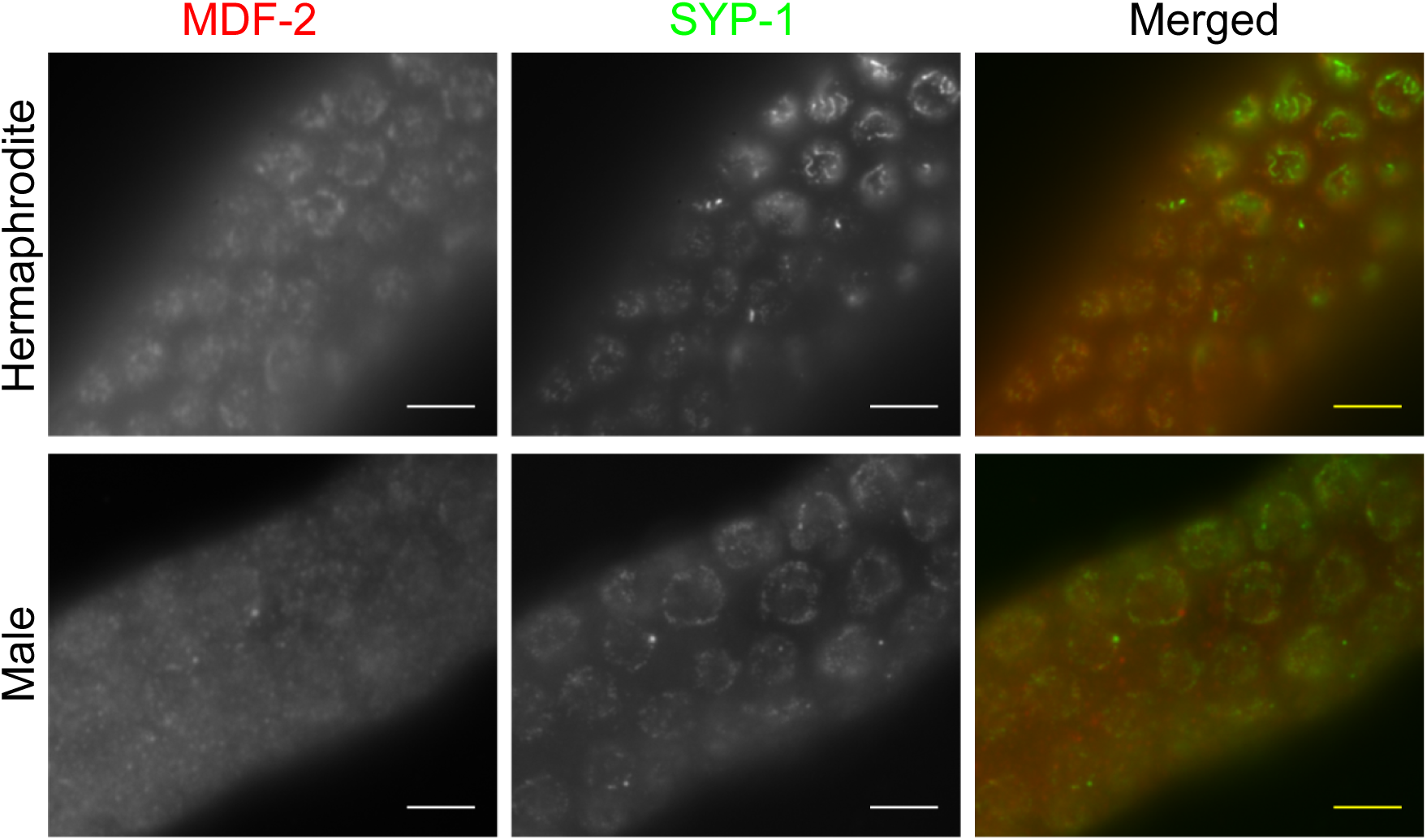
MDF-2 localization in early meiosis changes in males. (A) Whole mount fixed gonads of N2 young adults probed with anti SYP-1 (green) and anti MDF-2 (red) are shown with meiotic progression from left to right. MDF-2 localizes to the nuclear periphery during early meiosis before chromosomes begin synapsing in hermaphrodites (top row). However, males have more diffuse localization of MDF-2 before chromosomes begin synapsing (bottom row). Scale bars 5 μm.

To further investigate the MDF proteins in male vs female germlines, we next examined the rate of synapsis in *mdf* mutants. It was previously shown that mutants in the SAC (*mdf-1* and *bub-*3) have faster completion of synapsis. As *mdf-2* mutants have an extended mitotic region, and MDF-2 has differential localization in males, our analysis differed slightly from that previously published (Bohr *et al.* 2015). In this analysis, we measured the meiotic region according to where SYP proteins begin forming small foci before synapsis occurs and where HTP-3 starts localizing to chromatin (Fig. 8A). This was divided into five regions, and the percent of nuclei that had complete synapsis (all stretches of HTP-3 have co-localizing SYP-1 tracks) was measured (Fig 8A). We analyzed L4 hermaphrodites undergoing spermatogenesis, and there was no statistical significant difference in the percent synapsed in the first two zones of meiosis between N2 WT L4s and *mdf-2* L4s (Fig 8B). Unexpectedly, there was a significantly lower percentage of complete synapsis in the third and fourth zone in *mdf-2* mutants as compared to N2, indicating a minor full synapsis defect (p =0.0135, and p=0.049, respectively). However, a lack of difference in the first two zones of meiosis indicates that synapsis proceeds at the same rate in WT vs *mdf-2* spermatogenesis, further implying that MDF-2 does not function in the same way in male germ cells as in female germ cells. Additionally, all SAC proteins need to be present together for the SAC to be functional, and there is no difference between *mdf-1* L4 hermaphrodites and WT hermaphrodites. This further implies that there is a difference in the requirement for SAC function in male and female germ cells.

**Figure 8.**
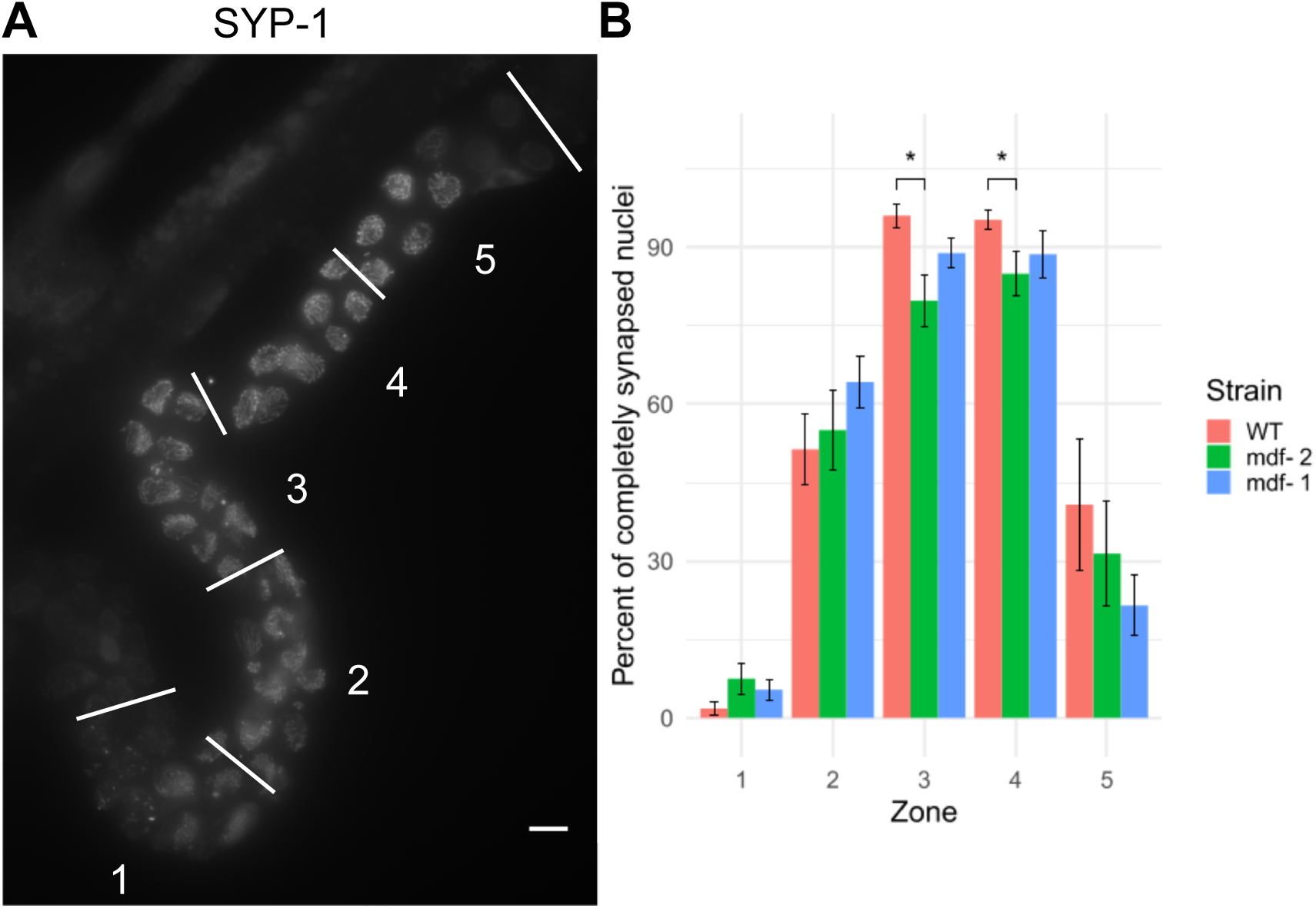
Loss of MDF-2 or MDF-1 does not increase rate of synapsis in male germ cells. (A) Whole mount fixed larval stage L4 hermaphrodite *mdf-2(av16) unc-17(e245)* IV gonad probed with anti SYP-1. Shown as an example of how the meiotic region of the gonad was measured starting at initial HTP-3 localization and SYP-1 foci until the end of pachytene; this region was divided into five equal length regions, and percent of nuclei synapsed (i.e. nuclei that all regions of HTP-3 tracks co-localized with SYP-1 tracks) was measured for each region. Scale bar 5 μm. Images stitched using Grid/Collection plugin for ImageJ developed by Preibisch et al., 2009. (B) *mdf-2* spermatogenic larval stage L4 hermaphrodites have the same rate of synapsis as WT L4 animals and *mdf-1* L4 animals in the first two zones. *mdf-2* L4 animals have a full synapsis defect in zones three and four, where they have a significantly lower rate of synapsis than WT. Error bars represent SEM.

## DISCUSSION

### A sex specific checkpoint at the initiation of synapsis

Chromosomes that are unable to complete synapsis are targeted for H3K9me2 enrichment during *C. elegans* meiosis (Bean *et al.* 2004). Surprisingly, whereas the complete absence of synapsis in male meiosis yields H3K9me2 enrichment on *all* chromosomes, complete asynapsis in female germ cells fails to trigger H3K9me2 enrichment on *any* chromosome. We have shown that in contrast to situations where the initiation of synapsis is inhibited, hermaphrodite meiotic nuclei that are able to partially initiate synapsis can still trigger enrichment of H3K9me2 in pachytene stage of hermaphrodite meiosis. It thus appears that the signal for H3K9me2 enrichment of unsynapsed chromatin in oogenesis requires at least partial synapsis initiation. In contrast, unsynapsed chromatin in males can be targeted for H3K9me2 enrichment, despite a complete lack of synapsis. In the absence of synapsis initiation, female oogenesis cannot proceed to the stage where H3K9me2 enrichment is activated, whereas in males this arrest does not occur, or if there is an arrest, it occurs after H3K9 methylation targeting. We propose that the differential regulation of H3K9me2 enrichment is a consequence of a checkpoint that is only present in oogenic meiosis.

Although it is a reliable marker of asynapsed chromatin in *C. elegans*, the exact purpose of H3K9me2 enrichment and its regulation is poorly understood, as it is not essential for meiotic progression. H3K9me2 enrichment on the unpaired X chromosome in XO (sex transformed) females has been shown to shield it from activating meiotic checkpoints, but detection of asynapsis of homologs is not affected by loss of H3K9me2 (Checchi and Engebrecht 2011). Mutations in *met-*2, the H3K9 methyltransferase responsible for germline H3K9me2, have only minor meiotic defects (Bessler *et al.* 2010). In addition, meiosis appears normal in *met-*2 mutant XO males, suggesting that the addition of H3K9me2 to unsynapsed chromatin is not essential for meiotic progression (Bessler *et al.* 2010). Despite its similarity to the Meiotic Silencing of Unpaired Chromatin (MSUC) process observed in other organisms, genes along unsynapsed chromosomes may not be repressed (Checchi and Engebrecht, 2011). It is thus not clear what consequences result from H3K9me2 enrichment, as it doesn’t appear to target genomic regions *de novo* on unsynapsed X chromosomes in *him-8* mutant hermaphrodites, nor are there obvious transcriptional changes (Guo *et al.* 2015). The enrichment for H3K9me2 in unsynapsed chromatin may be a simple consequence of defects in SC assembly and meiotic chromatin reorganization. Indeed, a previous study reported an increase in scattered foci of H3K9me2 in *syp-1* hermaphrodite germ cell chromatin (Lamelza and Bhalla 2012). Although we did not consistently observe this in multiple SC mutants, the difference in results could reflect a difference in the fixation procedures, which in turn could reflect some aspect of chromatin reorganization. It should also be noted that although we generally do not observe H3K9me2 enrichment coinciding where SC proteins have loaded, we do not always observe H3K9me2 where SC proteins are absent. Reorganization of meiotic chromatin may randomly expose genomic regions that are already heritably marked by H3K9me2 in germ cell chromatin, but are normally less accessible to MET-2 activity after axis maturation and SC assembly initiates. Alternatively, the increased enrichment of H3K9me2 may simply represent an alteration in chromatin structure that increases accessibility for the MET-2 methyltransferase only in males. However, it is unlikely that there is such a dramatic difference in chromatin accessibility between the sexes since *zim* mutants display enriched H3K9me2 on unsynapsed chromosomes in both sexes.

Increased methylation may be protective for the genome, as in the case of the un-partnered/unsynapsed X chromosome in XO hermaphrodites, and may mask its unsynapsed status and/or protect it from the activation of recombination checkpoints (Checchi and Engebrecht 2011). Importantly, MET-2 associates with the DNA repair protein SMRC-1 in mitotic and meiotic germ cells, and its activity is proposed to limit replication stress in germ cells (Yang *et al.* 2019). Loss of SMRC-1 leads to a generational depletion of H3K9me2 in germline chromatin, indicating it is required to heritably maintain MET-2-dependent H3K9me2. SMRC-1 is a worm homolog of SMARCAL1 and thus a member of the SWI/SNF family of ATP-dependent chromatin remodelers (Yang *et al.* 2019). The proposed checkpoint could act through SMRC-1 or other DNA damage sensing paths to recruit MET-2 to the exposed domains. Although there is no apparent difference in SMRC-1 presence or nuclear distribution in male versus hermaphrodites meiosis, its activity could be differentially regulated and this remains to be explored (Yang *et al.* 2019).

We therefore can only conclude that in female germ cells, the failure to initiate synapsis prevents meiotic progression to a stage in which H3K9me2 is targeted to unsynapsed chromatin, whatever the role of that targeting may be. In contrast, H3K9me2 targeting proceeds unimpeded in males, despite complete synaptic failure. Our interpretation of these results is that hermaphrodite meiosis requires passage of a post-pairing checkpoint that is absent in males. Interestingly, apoptosis doesn’t occur in *C. elegans* male germ cells (Gumienny *et al.* 1999; Gartner *et al.* 2000), yet even in hermaphrodite germlines activation of the newly proposed checkpoint may not induce apoptosis. Indeed, in *syp-1* hermaphrodites where the checkpoint appears to be activated, all apoptosis observed is accounted for by both the PCH-2 dependent checkpoint and by the DNA damage checkpoint (Bhalla and Dernburg 2005).

This newly proposed checkpoint is distinct from the PCH-2 dependent pachytene checkpoint as (a) we found no obvious difference in H3K9me2 enrichment (or lack thereof) when either HTP-3 or SYP-2 is depleted in the presence or absence of PCH-2, and (b) this proposed checkpoint takes place before the PCH-2 checkpoint. The PCH-2 dependent checkpoint senses the unsynapsed pairing centers remaining after other chromosomes have synapsed, and stalls meiosis in the pachytene stage (Bhalla and Dernburg 2005). Indeed, H3K9me2 enrichment on unsynapsed chromosomes is observed when either the DNA damage or pachytene checkpoints are activated, indicating that activation of these checkpoints does not preclude activation of MET-2 targeting to unsynapsed chromatin, suggesting they act downstream of H3K9me2 enrichment (Lamelza and Bhalla 2012). The proposed checkpoint senses failure to initiate *any* synapsis, leading to the stalling of meiosis in the leptotene/zygotene stage of prophase I. As we cannot currently separate the checkpoint from the absence of synapsis, it is unclear what the consequences of not having this proposed checkpoint are; it may even not be immediately essential for meiotic progression, as one of the only phenotypes measurable so far is initiation of H3K9me2 enrichment. However, as meiotic progression is stalled, a potential benefit of this proposed checkpoint would be that it delays further steps (e.g., redistribution of chromosomes throughout the nucleus from a clustered configuration) that could interfere with proper synapsis and allows further time for nuclei to finish pairing and begin synapsis if conditions are not ideal.

### Male synapsis is insensitive to high temperatures, unlike hermaphrodite synapsis

As has been shown previously, hermaphrodite SYP proteins form foci when grown at elevated temperatures (Bilgir *et al.* 2013). However, SYP aggregates do not form in either males or larval stage L4 hermaphrodites undergoing spermatogenesis when grown at an elevated temperature. This indicates that there is some process during synapsis initiation in female meiosis that is temperature sensitive, but this process is not apparent in spermatogenesis. The temperature sensitivity of this process in adult hermaphrodites could be the result of improper protein folding at this higher temperature, or this could be a mechanism to halt meiosis in an unfavorable environment. The temperature dependent component in female meiosis could be required for a signal once pairing has been established to initiate synapsis, or a protein that post-translationally modifies SYPs to allow assembly onto chromatin. Whatever the case, this process or protein is not necessary for synapsis initiation in spermatogenesis. *C. elegans* males typically result from non-disjunction of the X chromosome during hermaphrodite meiosis, and non-disjunction, and therefore males, occurs more frequently when hermaphrodite *C. elegans* are grown at higher temperatures. As males occur more frequently in a stressful condition like high temperature, and in higher temperatures the generation time for *C. elegans* compresses down to approximately 48 hours, it could be evolutionarily advantageous for male *C. elegans* to complete meiosis quickly and more robustly at elevated temperatures.

### Males mimic spindle assembly checkpoint mutants

While hermaphrodites require DLC-1 to synapse their chromosomes, males have grossly normal synapsis even with depletion of DLC-1. Males complete pairing and synapsis faster than hermaphrodites and even correctly pair their chromosomes without DLC-1. Male meiosis mimics spindle assembly checkpoint (SAC) mutant hermaphrodites, as these mutants also suppress the *dlc-1(RNAi)* phenotype and show an increased speed of synapsis completion. It has been hypothesized that the SAC proteins act to block or slow synapsis in early prophase I to allow more time for chromosomes to correctly pair (Bohr *et al.* 2015).

*C. elegans* males have an inherent problem in meiosis in that they have a single X chromosome with an exposed and unmatched pairing center. The spindle assembly checkpoint (SAC) complex is required for the pachytene checkpoint (Bohr *et al.* 2015), in which PCH-2 senses any unpaired pairing centers. This checkpoint could be triggered in every male meiotic nucleus because of the unpaired and unsynapsed X chromosome. Thus it is beneficial for males to not have a working pachytene checkpoint to keep meiosis from stalling because of their usual XO status. MDF-2, a component of the SAC, has a different localization in early male meiosis compared to females, as it is no longer concentrated near pairing chromosome ends and this could impede the function of the SAC. Without all of the SAC components, nuclei will be unable to sense that synapsis has not completed for all chromosome sets and therefore would not stall meiosis in males. Synapsis could complete more quickly in males without a fully functioning SAC complex to delay this step. While males do not undergo meiotic apoptosis in response to checkpoint activation, they still stall meiosis in response to DNA damage checkpoint activation (Gumienny *et al.* 1999; Gartner *et al.* 2000; Jaramillo-lambert *et al.* 2010). Therefore dismantling checkpoints that will unnecessarily hinder all meiotic progression both speeds up meiosis and conserves energy for male *C. elegans*, allowing them to produce more sperm and hence more progeny. Males complete prophase I two and a half to three times faster than hermaphrodites (Jaramillo-Lambert *et al.* 2007); the absence of a step designed to slow meiotic progression likely contributes to the accelerated speed of meiosis observed in male *C. elegans*.

### Parallels in *C. elegans* meiosis and mammalian meiosis

Mammals, like *C. elegans*, have differences in phenotypes related to SAC function in males vs females. Male mammals have a more stringent SAC response to problems in meiosis, such as to Robertsonian fusions, including arresting meiosis at meiotic metaphase (Eaker *et al.* 2001), while females do not show evidence for metaphase arrest (Chmatal *et al.* 2014). In mammals, the reduction in stringency in females appears to be linked to the large size of oocytes, with the resulting dilution of the SAC proteins in the larger volume restricting the ability of the SAC to prevent the transition to anaphase (Kyogoku and Kitajima 2017). Similarly, lower levels of expression and/or a change of localization of any of these SAC proteins in *C. elegans* males could lead to a lower local concentration of these proteins as in mammalian oocytes, and therefore to less activity.

Spermatogenesis in mammals appears to be more greatly affected by several defects in meiotic processes, which usually result in meiotic arrest at early stages of prophase I (reviewed in (Morelli and Cohen 2005)), and infertility or greatly reduced fertility in males. However, these same mutations in oogenesis in mammals results in arrest at a later stage of prophase I, and several mutations still result in fertile females. This general reduction of stringency for oogenesis checkpoints in mammals is opposite to what we have observed in *C. elegans* spermatogenesis. We have shown that male *C. elegans* pair and synapse their chromosomes faster, do not require DLC-1 to synapse or pair their chromosomes, and are able to synapse chromosomes when raised at high temperatures, all or any of which may or may not be linked to a novel checkpoint that senses the initiation of synapsis. All of these differences may ultimately result from sex chromosome evolution in a hermaphroditic species with the stress-induced generation of XO males. *C. elegans* males therefore may be able to give further insight into other ways that meiotic checkpoints can be escaped by one sex but not the other, and how these differences evolve.

## Supplemental Figures

**Figure S1.**
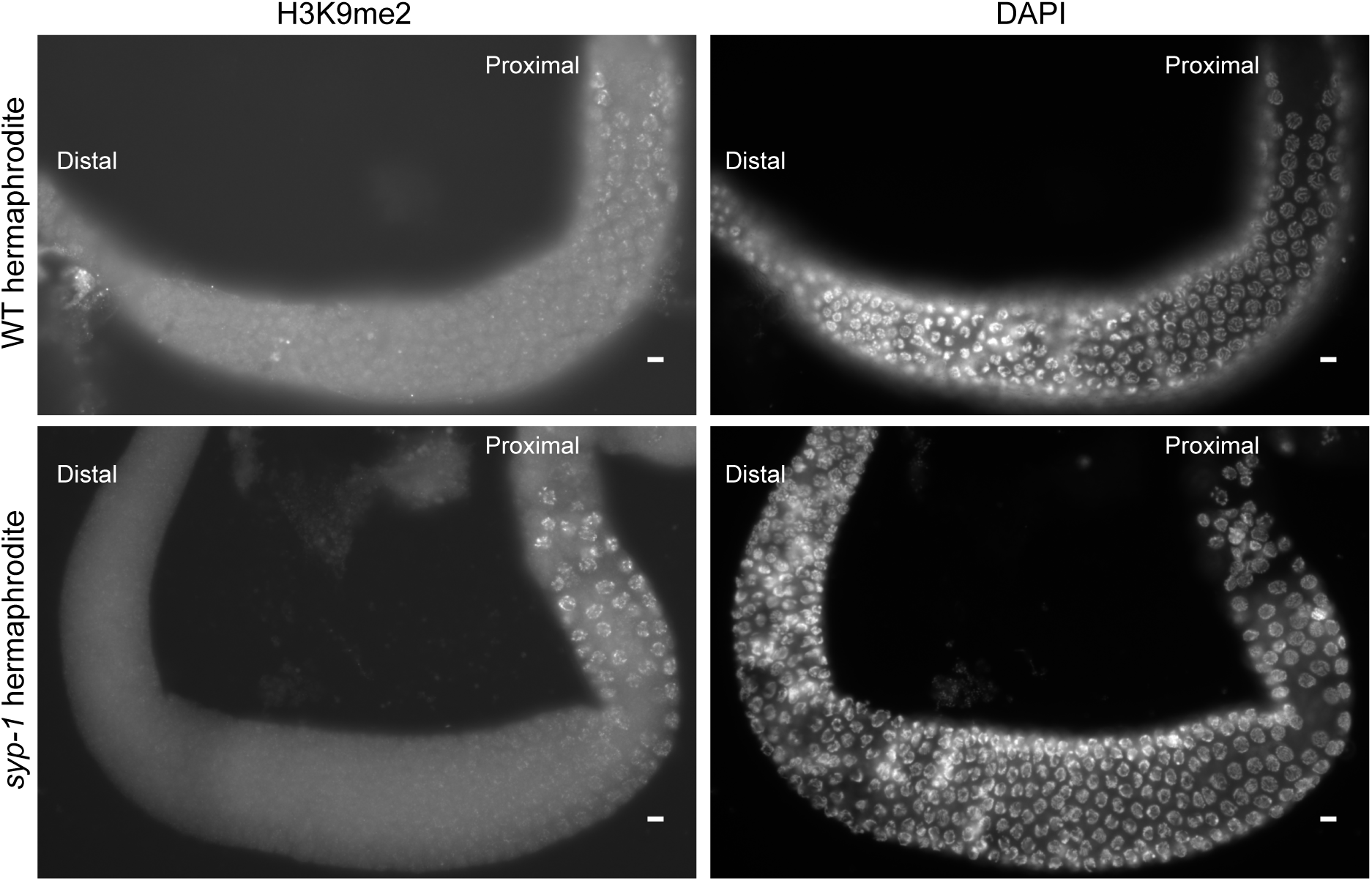
Both WT and *syp-1* hermaphrodites have widespread enrichment of H3K9me2 in late pachytene, early diplotene. Whole mount fixed gonads probed with DAPI and anti H3K9me2, with meiotic progression from left to right. WT hermpahrodites have low levels of H3K9me2 throughout prophase I, until late pachytene when there is widespread enrichment (top row). *syp-1* hermaphrodites also show low levels of H3K9me2 until late pachytene, when they also display widespread enrichment (bottom row). Both images were cropped and used for Figure 1. Scale bars 5 μm.

**Figure S2.**
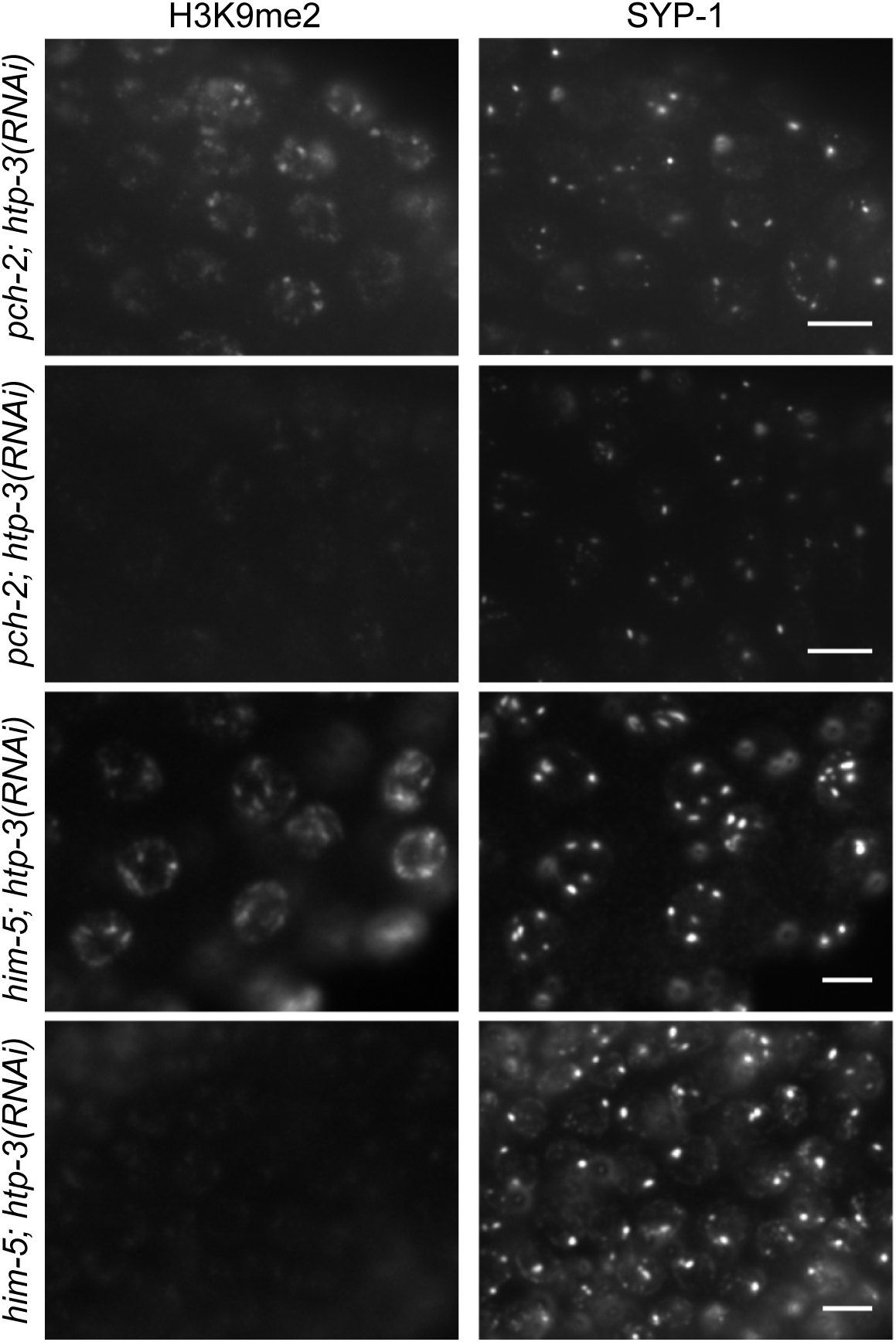
H3K9me2 stays low with depletion of HTP-3 with occasional exceptions in hermaphrodties. Whole mount fixed gonads of either *pch-2* or *him-5* hermaphrodites probed with anti-SYP-1 and anti-H3K9me2 grown with *htp-3* RNAi. *pch-2* hermaphrodites with HTP-3 knockdown display SYP-1 polycomplexes, and occasional widespread enrichment of H3K9me2 (top row), while most gonads show low H3K9me2 when SYP-1 is in polycomplexes (second row). However, this also occurs with knockdown of HTP-3 in hermaphrodites with normal PCH-2 function: occasional widespread enrichment of H3K9me2 with SYP-1 in polycomplexes (third row), with most gonads displaying low H3K9me2 when SYP-1 is in polycomplexes (bottom row). Scale bars 5 μm.

## Notes

#### Summary of Updates

New title and abstract

